## Supplementary material for "Pixel: a content management platform for quantitative omics data": Table 1

| **Omics Unit** | **Description** | **A** | **B** | **A (QS)** | **B (QS)** |
| --- | --- | --- | --- | --- | --- |
| 1. CAGL0F04807g | Ortholog(s) have role in **pathogenesis** and cell surface, hyphal cell wall, integral component of mitochondrial outer membrane, plasma membrane localization | 1,09 | 1,81 | 2,23E-19 | 7,31E-05 |
| 2. CAGL0F06457g | Ortholog(s) have role in fungal-type cell wall organization or biogenesis, mitochondrial outer membrane translocase complex assembly, **pathogenesis**, phospholipid transport, protein import into mitochondrial outer membrane | 0,30 | 0,19 | 4,14E-02 | 2,65E-01 |
| 3. CAGL0I02970g | Ortholog(s) have delta14-sterol reductase activity and role in cellular response to drug, ergosterol biosynthetic process, filamentous growth of a population of unicellular organisms in response to biotic stimulus, **pathogenesis** | 0,90 | -2,64 | 4,65E-16 | 2,19E-05 |
| 4. CAGL0I10516g | Ortholog(s) have role in fungal-type cell wall organization, pathogenesis and cytoplasm, eisosome, integral component of plasma membrane, membrane raft localization | 1,50 | 0,57 | 8,29E-60 | 1,16E-02 |
| 5. CAGL0L08448g | Ortholog(s) have role in actin cytoskeleton organization, eisosome assembly, negative regulation of protein phosphorylation, negative regulation of sphingolipid biosynthetic process and **pathogenesis**, more | 1,67 | -0,57 | 1,77E-75 | 7,04E-03 |
